# Measuring phase-amplitude coupling opposition in neurophysiological signals with the Mean Opposition Vector Index (MOVI)

**DOI:** 10.1101/2023.08.04.551929

**Authors:** Ludovico Saint-Amour di Chanaz, Alexis Pérez-Bellido, Lluís Fuentemilla

## Abstract

**Background:** The phase–amplitude coupling (PAC) opposition between distinct neural oscillations is critical to understanding brain functions. Available methods to assess phase-preference differences between conditions rely on density of occurrences. Other methods like the Kullback-Leibler Divergence (DKL) assess the distance between two conditions by transforming neurophysiological data into probabilistic distributions of phase-preference and assessing the distance between them. However, these methods have limitations such as susceptibility to noise and bias.

**New Method:** We propose the “Mean Opposition Vector Index” (MOVI), a parameter-free, data-driven algorithm for unbiased estimation of PAC opposition. MOVI establishes a unified framework that integrates the strength of PAC to account for reliable unimodal differences in phase-specific amplitude coupling between neurophysiological datasets.

**Results:** We found that MOVI accurately detected phase opposition, was resistant to noise and gave consistent results with low or asymmetrical number of trials, therefore in conditions more similar to experimental studies.

**Comparison with existing methods:** MOVI outperformed Jensen-Shannon Divergence (JSD), an adaptation of the DKL, in terms of sensitivity, specificity, and accuracy to detect phase opposition.

**Conclusions:** MOVI provides a novel and useful approach to study of phase-preference opposition in neurophysiological datasets.

## 1. Introduction

Neurophysiological data exhibits oscillations of different frequencies, which may co-occur and also interact with one another (Hyafil, Giraud, Fontolan, & Gutkin, 2015; Jensen & Colgin, 2007; Tort, Komorowski, Eichenbaum, & Kopell, 2010). Among the various forms of coordination, phase-amplitude coupling (PAC) stands out as the most widely acknowledged as it can be detected throughout the brain in animal studies (Jensen & Lisman, 1996; Pavlides, Greenstein, Grudman, & Winson, 1988; Tort et al., 2008) and human brain recordings, using techniques such as electroencephalography (EEG) (Axmacher et al., 2010; Canolty et al., 2006; Tort, Komorowski, Manns, Kopell, & Eichenbaum, 2009), magnetoencephalography (MEG) (Fuentemilla, Penny, Cashdollar, Bunzeck, & Duzel, 2010) and local field potential (LFP) recordings (Rutishauser, Ross, Mamelak, & Schuman, 2010; Saint Amour di Chanaz et al., 2023). PAC involves the interaction between the phase (the timing) of low-frequency brain oscillations and the amplitude (the strength) of higher-frequency oscillations and outlines that information coding in the brain involves more than just individual neuron action potential firing, but the brain’s extracellular field potential oscillations serve as temporal reference to aid efficient information processing by the spiking activity of the neural ensembles.

The best studied-example is the coupling of the ongoing theta (3-8Hz) and the amplitude of gamma oscillations (>30Hz) (Bragin et al., 1995; Buzsaki, 2002; Tort et al., 2008) Theta-gamma PAC has been reported in a variety of species, brain regions and experimental conditions (Axmacher et al., 2010; Giraud & Poeppel, 2012; J. E. Lisman & Jensen, 2013), and theoretical work has suggested hippocampal theta-gamma PAC supports a neural coding regime to support mnemonic operations (Colgin, 2015) and spatial navigation (Huxter, Senior, Allen, & Csicsvari, 2008; Jensen & Lisman, 1996; O’Keefe & Burgess, 2005; Tsodyks, Skaggs, Sejnowski, & McNaughton, 1996). The importance of theta-gamma PAC is that the mechanisms behind it provide valuable insights into how the brain processes and represents information. For example, influential computational models, supported by empirical findings in rodents (Bragin et al., 1995; Colgin et al., 2009) and in human (Kunz et al., 2019; Pacheco Estefan et al., 2021) posit that theta-nested gamma oscillations may allow the hippocampal network to temporally organize sequences of events within each theta cycle (J. Lisman, 2005; J. E. Lisman & Idiart, 1995; J. E. Lisman & Jensen, 2013; O’Keefe & Recce, 1993) Other models, also supported by empirical data in rodents (Bieri, Bobbitt, & Colgin, 2014; Colgin et al., 2009; Douchamps, Jeewajee, Blundell, Burgess, & Lever, 2013; Fernandez-Ruiz et al., 2017; Lever et al., 2010; Manns, Zilli, Ong, Hasselmo, & Eichenbaum, 2007; Pavlides et al., 1988; Poulter, Lee, Dachtler, Wills, & Lever, 2021) and in humans (Saint Amour di Chanaz et al., 2023), propose that gamma couples to opposed-phase states of the ongoing hippocampal theta rhythm activity during memory encoding and during retrieval (Hasselmo, Bodelon, & Wyble, 2002). Together, these findings underscore that neural oscillations have broader implications than just PAC alone. Taking the phase preference of PAC into account is critical, as it significantly influences the organization and processing of mnemonic information in the brain. Therefore, it becomes essential to incorporate analytical tools that can effectively characterize differences in phase preference within PAC across neural states.

There are several quantitative approaches to identify phase preference within PAC and how phase preference within PAC differs or opposes across neural states or experimental conditions. Some of these methods rely on the existence of PAC and aim at detecting phase-preference differences between two conditions. Analytically, these methods, such as the Inter-Trial Phase Coherence (ITPC);(VanRullen, 2016), are used to estimate average angular degree of incidences in which each trial is summarized into a single vector with a phase preference and then the angles of the vectors are compared to a uniform polar distribution or to another subset of trials. These methods rely, therefore, on density of occurrences. An important limitation in the context of detecting phase preference within PAC, however, is that these approaches do not consider coupling strength of angle preference as each trial, or observation is summarized into a unit-normalized vector. Thus, while these methods are suitable to quantify phase clustering across trials or observations, they undermine the possibility that phase preference could be brought by weak or even non-meaningful amplitude modulations. In these cases, these indexes could still provide non-random phase clustering values but that would compromise the interpretability of the underlying neural mechanisms.

Neurophysiological data can also be analysed using PAC methods implemented separately into equal phase bins. In PAC analysis, phase bins are used to divide the phase values of low-frequency oscillations (e.g., theta) into equal intervals or segments. These intervals represent different phase angles that cover the entire 360-degree range of a circular distribution. For each bin, the mean amplitude is calculated, resulting in a distribution of amplitude per phase. This process combines the time series of phase and amplitude into a single distribution of amplitude per phase, capturing both components without losing information about the variability of amplitude phase-preference. The method involves taking the time-series of the filtered amplitude and phase, identifying time-points corresponding to specific phase ranges based on the number of bins, and then averaging the amplitude time-points to obtain a single average amplitude measure for each phase bracket. The distributions obtained from the data can be studied using different methods, such as the Kullback-Leibler Divergence (DKL). DKL is a commonly used method in fields like genetics (Akhter et al., 2017) and engineering (Ji et al., 2022) to measure the distance between two probabilistic distributions. However, a limitation of DKL is that it is not symmetric in comparing distributions (Nielsen, 2019). This means that the distance between distribution A to B is not the same as the distance from distribution B to A, which can lead to asymmetrical results when comparing distributions using DKL.

The Jensen-Shannon Divergence (JSD) overcomes the asymmetry limitation of the DKL by providing a symmetrical measure (Nielsen, 2019). It calculates the distance between the mean distribution of two datasets (A and B) and then compares each dataset to the mean distribution, resulting in a symmetrisation of the DKL. However, JSD uses a logarithmic transformation, making it more suitable for detecting average differences between distributions rather than those driven by a single-phase preference. Another shortcoming of JSD is it has the capability to detect multimodal differences in phase preference, which can be problematic. If one or both distributions exhibit multiple peaks of phase preference (multimodal), JSD might identify a difference even in situations where it may not reflect a clear phase preference. For example, if high-frequency activity is coupled to 8Hz at one specific phase (unimodal), JSD may erroneously indicate a bimodal coupling at 4Hz with two peaks of phase preference. This sensitivity to multimodal distributions is not ideal when assessing phase-preference differences, as it can lead to misinterpretations and biases, especially towards lower modulating frequencies.

To address the limitations of the JSD in assessing uniform and unimodal opposition in phase preference, we developed a new algorithm that uses the Mean Vector Length (MVL) measure (Canolty et al., 2006) and creates an alternative distribution from the difference of the two distributions being compared. By applying the MVL to this new distribution, we can detect a uniform and unimodal opposition in phase preference while considering phase-amplitude coupling strength. We refer to this index as Mean Opposition Vector Length (MOVI), and we will detail below its premises, assumptions, and usefulness when working with neurophysiological data. In this study, we will compare MOVI against JSD, that encompasses most of the criteria we wanted to assess phase-opposition by considering coupling strength, not risking of having abnormally high or low numerical values like Jeffrey’s Divergence, and symmetry in the sense that the distance between A and B would be the same as the distance between B and A. We will contrast their differences in terms of sensitivity and specificity (Trevethan, 2017), calculate their predictive value, and assess accuracy and Matthews Correlation Coefficient (MCC) (Chicco, Totsch, & Jurman, 2021) on simulated data, where we will manipulate different variables and parameters that can be found in neurophysiological data such as PAC strength, noise, opposition angle, number of trials, and inter-trial phase coherence. Matlab scripts to produce the simulations and perform the analyses described in this paper are available at github: https://github.com/DMFresearchlab/MOVICode

## 2. Material and Methods

### 2.1. Synthetic data generation

We simulated datasets containing two phase-coupled trial distributions. For both distributions we simulated a time series of coupled low and high frequencies. The frequency sampling of these time-series was of 1000 Hz and we simulated 2500 time points, or the equivalent of 2.5 sec epochs. When generating our datasets we simulated each trial independently using the formulas provided by (Tort et al., 2010) and adding variable parameters to control for coupling angle for each distribution. We simulated a baseline of 50 trials, and each trial was composed by a pair of frequencies, a frequency for phase (*f*(*p*)) and a frequency for amplitude (*f*(*a*)). We also took into account the filtering parameters, and hence simulated the central frequencies of interest (f) as well as the low and high limits of the frequency window *f*(*a*) ± *f*(*a*) ∗ 0.35 for amplitude and *f*(*p*) ± *f*(*p*) ∗ 0.2 for phase. All trials were simulated in the following way:

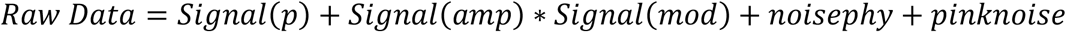

Where

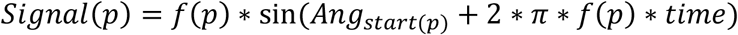

and

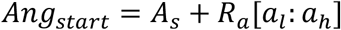

The starting angle of each frequency was determined by the sum of a fixed starting angle *A*_*s*_ and a random angle *R*_*a*_ between a lower limit *a*_*l*_ and a higher limit *a*_*h*_. The random angle made each trial slightly different within each condition but within a fixed angle range. We wilfully introduced this noise to have a similar phase preference across trials and to try to replicate real neurophysiological data where different trials do not have the exact same characteristics. It also helped to ensure that each trial was different to justify correction by surrogate trials. Without the random angle all trials would have been identical and averaging over trials would be the same as having one single trial. In this case the correction by surrogate trials would also have been unnecessary.

*Signal*(*amp*) was generated in the same way as *Signal*(*p*) but with a beginning angle specific to amplitude *Ang*_*start*(*amp*)_.

*Signal*(*mod*)) is the modulatory signal of amplitude responsible for cross-frequency coupling mechanisms and it was defined in the following way:

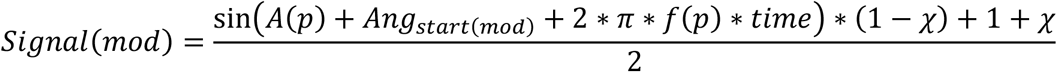

where, *A*(*p*) is the fixed starting angle for phase to ensure that the modulation of the signal is locked to the phase for each trial. *Ang*_*start*(*mod*)_ is the starting angle specific to modulation and relative to the phase angle where the modulation can happen in phase or out of phase with the low frequency. *χ* is a coupling factor between 0 and 1, where 1 is no coupling and 0 is perfect coupling. The modulatory *sin* wave always ranges between 0 and 1 and by multiplying the amplitude time series with the modulatory time series we ensure that we obtain a high frequency coupled to the frequency of the modulatory wave.

Noise adding uncoupled low and high frequencies to simulate as closely as possible noise that is likely to be found in real neurophysiological data. First, we ensured that the raw signal was slightly different for every trial by adding a low-frequency wave with random starting angle, and a non-modulated high frequency wave. Then, pink noise (van Driel, Cox, & Cohen, 2015) was added with an amplitude of the noise signal being equivalent to the amplitude of the coupled high frequency signal. We then computed MOVI and JSD on simulated data with these characteristics 100 times, which emulated an experiment with 100 synthetic participants, each with the same number of trials and same settings. Knowing beforehand the degree of coupling and opposition allowed us to know the ground truth and benchmark the different methods used to assess for opposition. Since within the settings there was randomness added with random starting angle within a fixed angle window or with physiological-like noise it ensured that each synthetic participant was different. This allowed us to have a distribution of the MOVI and JSD scores to assess also how much variability there was in the indexes and if any factor or setting particularly influenced sensitivity or specificity. When the degree of angle opposition was not parametrically manipulated, the two datasets had an opposition of 0° (non-opposed) or 180° (completely opposed). When not manipulated, PAC strength of both distributions was average and fixed, both distributions had 50 trials and each trial had a randomness of inter trial coherence of 180°, meaning that all trials had a phase preference that was within ±90 degrees from the manually determined angle of coupling.

### 2.2. Pre-processing of the signal

Each trial of raw data was filtered using the eegfilt function of EEGlab (Delorme & Makeig, 2004). The window of the frequency of interest (f) was determined as well as *f*(*a*) ± *f*(*a*) ∗ 0.35 for amplitude and *f*(*p*) ± *f*(*p*) ∗ 0.2 for phase. Filtering was done on longer epochs that were then cut to fit the desired number of time points, removing any filtering artefacts. The filtering window increased with frequencies for both phase and amplitude extraction in order to avoid biases towards lower modulating frequencies and ensuring that the window of filtered high frequencies was always superior to twice the phase of locking: Δ_*amp*_ > 2 ∗ *phase* (Aru et al., 2015). Then we extracted angles of phases for low-frequencies, and we extracted the absolute value of the square of the Hilbert envelope of amplitudes for high frequencies. Epochs were then binned in 18 equal bins of 20° (Tort et al., 2010). Each trial was then converted to a probabilistic distribution where sum = 1 by dividing the amplitude of each bin by the sum of all bins. Subsequently, we used these bins to calculate JSD and MOVI between different trial conditions.

### 2.3. JSD calculation

JSD requires the extraction of the DKL index, which is described as follows:

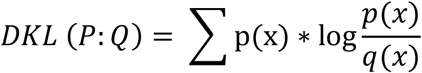

where *p*(*x*) is the value of each bin of distribution *P, q*(*x*) is the value of each bin of the distribution *Q*. This measure, however, is asymmetrical, in the sense that *DKL* (*P: Q*) is different than *D*KL (*P: Q*). This is problematic in the sense that to assess distance or opposition of angles it is important that the choice of comparison order (distance from *P* to *Q* or from *Q* to *P*) does not alter the results. It is possible to make DKL symmetrical considering Jeffrey’s Divergence (Nielsen, 2019) with the following formula:

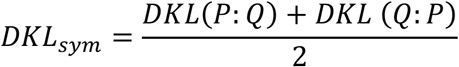

This symmetrisation, however, has been shown to have numerical issues where resulting values were either too high or too low (Nielsen, 2019) and an alternative approach to make the measure symmetrical was developed using the Jensen-Shannon Divergence. The Jansen-Shannon Divergence uses DKL formulas to estimate the distance of each distribution to the average between the two distributions in the following way:

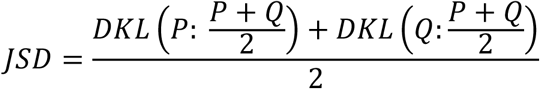

Where 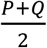 is the average distribution of *P* and *Q*. This distance can be interpreted as the total divergence from the average distribution. The important property of the Jensen-Shannon divergence compared to the symmetrical DKL is that this distance score is always bounded 0 ≤ *JSD* ≤ *log*2 and reduces biased opposition estimated given by the symmetrical DKL.

### 2.4. MOVI calculation

To calculate MOVI, like in JSD, we used the MVL on an alternate distribution made with the distributions A and B as explained below. First, we averaged the trial distributions of A and B to have a single distribution that was descriptive of common PAC effects and that shared a common direction. By doing so, if all trials exhibited strong PAC but each of them had their amplitude coupled to a different phase of low frequencies, the average distribution was flat, and no PAC would be detected. Measuring PAC on the average distributions of instead of doing it at the trial level allows for a measure of phase amplitude coupling across trials that share a direction preference and prevent sensitivity to spurious trial-level PAC that might not systematically rely on the same phase preference. In sum, averaging over trials is important because MOVI aims to detect phase-preference opposition between two conditions, and we need common directional effects of two distributions to assess opposition. For MOVI to be significant there needs to be a consistency in phase preference across trials (ITC) but also a consistency of coupling strength (PAC), and that this be the case for both compared distributions. If all trials are strongly coupled but each to a different phase in on or both distributions, MOVI will not be significant. Similarly, if all trials have a consistent phase preference in both distributions but a weak coupling, MOVI will also not detect opposition. We performed these steps for two distributions that had either similar or opposed angles of phase preference. Both averaged distributions were then converted to a probabilistic distribution (sum = 1) before comparing them with the different techniques.

To compute MOVI, we took the two distributions A and B obtained after the implementation of pre-processing stage and created an alternative distribution (*D*_*alt*_)of amplitude per phase bin based on the difference of the two original distributions with the following formula:

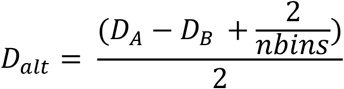

where *D*_*A*_ is the distribution of A trials, *D*_*B*_ is distribution of B trials and *nbins* is the number of bins (18 in this case).

We then obtained the MVL measure (Canolty et al., 2006; Cohen, 2008) to the alternative distribution *D*_*alt*_ to assess the strength of opposition between the two distributions averaged over trials.

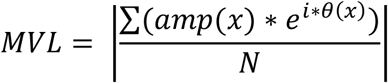

where *amp*(*x*) is the amplitude of each phase bin (*x*), *θ*(*x*) being the phase of each bin (*x*) and *N* the number of bins. This process vectorised each phase bin and extracted the absolute value of the mean vector, which gives the vector length. The longer the vector, the higher the coupling, or in the case of MOVI, the opposition.

We added twice the value of the uniform distribution 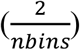 to avoid having negative values and facilitate the normalisation of the distribution into a probabilistic one (sum = 1). Then, we divided everything by 2 to have an alternative distribution *D*_*alt*_ that had a peak to through difference magnitude equivalent to either distribution. In the case of opposition in phase preference, *D*_*alt*_ would have a non-uniformity score equal to the sum of the MVL of both distributions, by dividing the alternative distribution by 2 we ensure that the MVL magnitude of this alternative distribution is instead between the MVL of both distributions. This way, we ensure that PAC strength is taken into consideration, and we lower the risk of having false positives by not increasing the peak to through difference.

*We* hypothesized that if A and B distributions have opposed phase preferences, the alternative distribution would be unidirectional and non-uniform and would exhibit a strong MVL (**Figure 1A**). On the other hand, if distributions A and B have the same phase preference, the alternative distribution would be flat and the MVL would be 0 (**Figure 1B**). In the case of both A and B distributions showing a weak PAC, even in conditions where the phase preferences were opposed between two, MOVI would be small. MOVI, therefore, ensures that phase opposition would be statistically significant only when the two distributions showed a solid phase opposition and a strong PAC.

**Figure 1.**
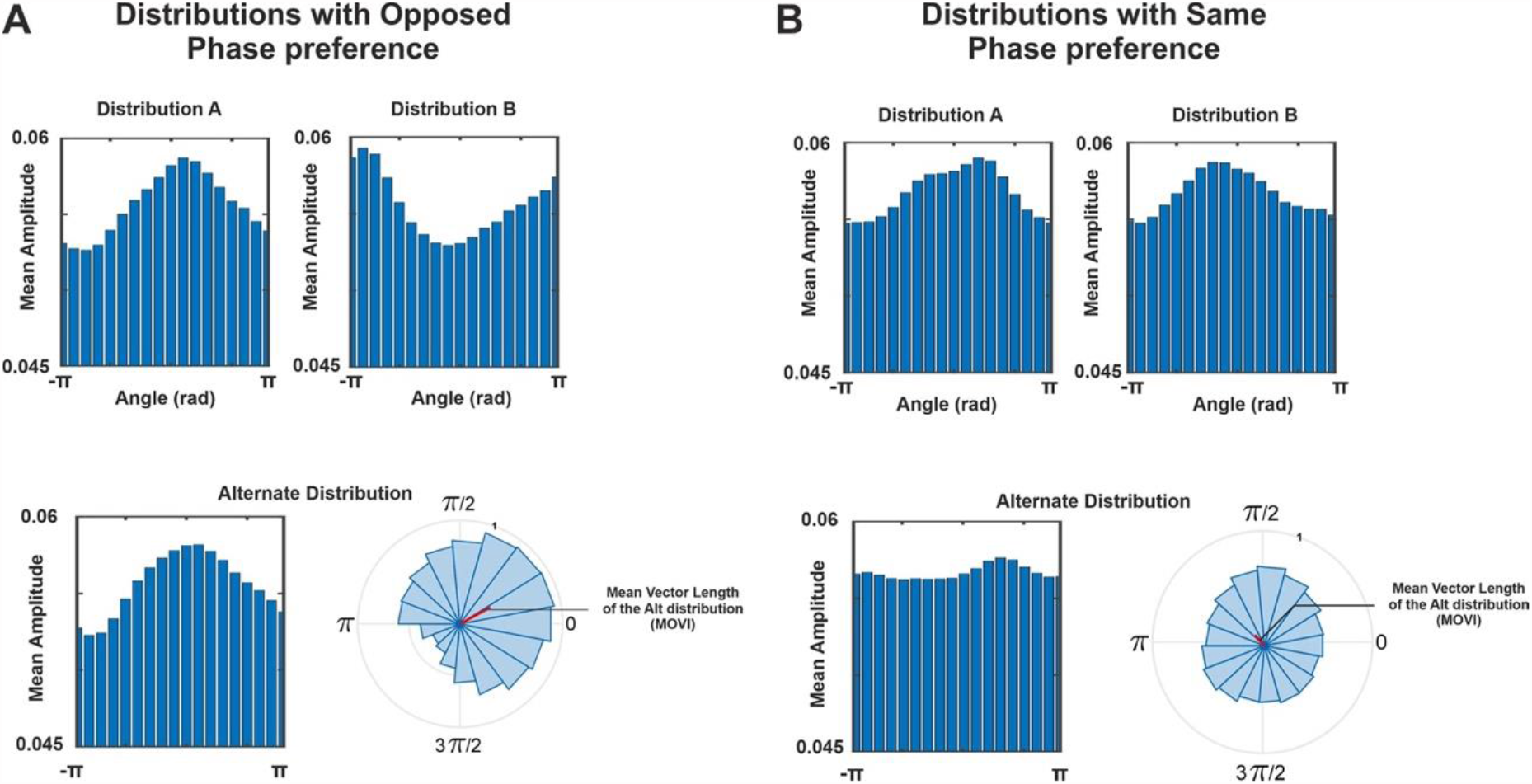
Schematic representation of MOVI. (**A**) Distributions A and B exhibit high PAC and opposed phase preference. The alternate distribution exhibits a strong preference. The high MVL value of the alternative distribution indicates opposition of phase preference between distributions A and B. (**B**) Distributions A and B exhibit high PAC but the same phase preference. The alternate distribution thus shows no phase preference and is flat. MOVI has a low value and indicates that distributions A and B are not opposed.

### 2.5. Statistical analyses

The statistical assessment of MOVI and JSD involves constructing a null distribution employing surrogate trials, which are alternative trials distributions with shared variance. These surrogate trials are utilized to consider any common effects observed in the experimental data. This non-parametric approach results in surrogate trials having a MOVI or JSD score that is normally distributed around a mean. The assessment of significance of the experimental value is done by comparing that experimental value to the distribution of the surrogate values.

For each comparison of A and B using either MOVI or JSD, where a comparison is made for every specific set of variables (angle, PAC strength, frequency, ITC, etc) we computed 1000 surrogate trials for each condition A and B. Surrogate trials may be calculated in two ways. First, label shuffling, which consists in shuffling the labels of amplitude trials before binning. Label shuffling creates a surrogate distribution that corrects for neural responses not necessarily related to phase-amplitude coupling (e.g., ERPs or phase resetting) but loses individual coupling that is more specific to each trial. Second, time-shuffling, which involves the random division of each amplitude trial into two segments, with the subsequent inversion of the order of the time-series chunks within the trial. This way a random coupling would be created. However, this method is less conservative, as it does not account for effects that might be driven by an ERP or a phase reset mechanisms. In general, both of these surrogate trial creation methods yield highly comparable outcomes, and the decision to prefer one over the other primarily depends on the extent to which one desires or requires control over ERP or phase-reset driven effects. Although we did not expect to observe results driven by phase-reset in synthetic data, we here used the label-shuffling method as this would control for these effects on real neurophysiological data. Therefore, we shuffled the labels of amplitude trials and randomly matched them with phase trials before binning and averaging the binned distributions over trials. Once an average null distribution was generated for each of the two experimental conditions, we calculated the MOVI and JSD index in the same way as for the non-shuffled trials.

When conducting a test for MOVI or JSD statistical significance for a single pair of frequencies, we compared the experimental value of MOVI or JSD to the respective distributions of surrogate scores. We considered a test to be significant if the experimental score was higher than 95% of the surrogate scores. The p-value was calculated as 1 minus the proportion of surrogates lower than the experimental value. We computed this for 100 artificial subjects to see consistency of testing and false alarm or false negative rate when varying parameters in the simulation like PAC strength, or opposition angle.

### 2.6. MOVI and JSD comparison

Using synthetic data is useful to quantify the efficacy of MOVI against the JSD approach because it allows us to predefine the ground truth (i.e., opposition or not) and objectively estimate the difference in sensitivity (proportion of positive tests for truly opposed distributions) and specificity (percentage of negative tests for non-opposed distributions) for both methods of analysis as a function of different parameter manipulations. We compared MOVI and JSD by calculating the proportion of hits (significant opposition when actual opposition exists) and the proportion of false alarms (significant opposition when no actual opposition exists), as a function of the manipulation of i) PAC strength of both or either distribution, ii) angle of opposition between the two distributions, iii) number of trials, iv) inter trial coherence, and v) noise strength.

Results are displayed using a 2D matrix that shows the proportion of how often a test is significant given a specific set of variables for both MOVI and JSD. To estimate the sensitivity of a measure, we quantified how well MOVI or JSD identified true positives (characterizing type I error). To quantify the specificity, we quantified how well MOVI or JSD identified true negatives, or the absence of false positives (characterizing type II error). In other words, when we observe false positives, specificity decreases and type II error increases. The hits and false alarms matrices were then compared to see if there was a significant difference in sensitivity and specificity between both opposition measures using a binomial test. P-values were obtained for each combination of variables (PAC, Angle, number of trials, etc.). A p < 0.05 was used to threshold statistical significance. However, we used a Bonferroni correction to compensate for multiple comparisons, and showed only the values that survived the correction. In the results section we report the minimum and maximum significant t and p-values in each matrix.

We also computed the average Positive Predictive Value (PPV) of the matrices for each index and generated a PPV and a Negative Predictibe Value (NPV) matrix for each index and taking as observation the individual voxels of each matrix. We then compared these values with a t-test with the degrees of freedom being equal to the number of voxels per matrix -1 (in this case df = 24). PPV is the ratio of true positives over the number of total positive tests. It ensures that an increase in sensitivity in a test that likely induces more false positives does not result in a too steep decrease of specificity (Trevethan, 2017). PPV is the probability of having a real effect if the test is positive. For example, a PPV of 0.97 states that a positive test has 97% chances of being a true positive rather than a false positive. PPV is linked with the false discovery rate (FDR) as FDR = 1-PPV and PPV = 1-FDR, where FDR is the ratio of true positives over all positive tests.

The PPV is calculated as:

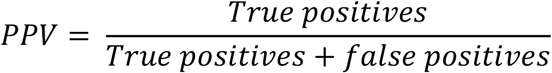

The average PPV was calculated for both each value of the 2D result matrices that resulted from MOVI and JSD approach and then compared with a repeated measures t-test. Similarly, we calculated the NPV that assesses the probability of an effect being truly negative given a negative test. NPV is linked with the False Omission Rate (FOR) as NPV = 1-FOR and FOR = 1-NPV, where FOR is the ratio of False Negatives over all negative tests. To assess how a test correctly identifies positive and negative outcomes, we also calculated the Accuracy as:

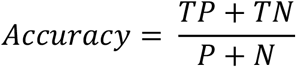

where TP are true positives, TN are true negatives, P are all positive tests and N are all negative tests. Finally we can assess an index performance with Matthews Correlation Coefficient (MCC) (Chicco et al., 2021) that has a value between -1 and 1. Since MCC is a measure that encompasses performance and bias, a value of 1 would mean that the test detects all the effects with no bias, while a value of -1 would be a test that does not detect effects at all and has only biases or false discoveries. MCC is calculated as follows:

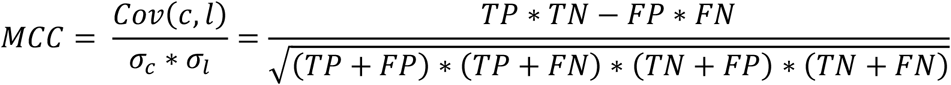

where *Cov*(*c, l*) is the covariance of true classes *c* and predicted labels *l*, and *σ*_*c*_ and *σ*_*l*_ are their standard deviations, respectively. The MCC is related to the other measures we explained below as:

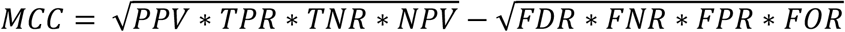

where TPR is the true positive rate, TNR true negative rate, FNR is the false negative rate, FPR false positive rate (Chicco et al., 2021).

## 3. Results

We simulated different conditions that typically happen and constrain the capacity to detect phase opposition by inflating Type I or Type II error rates in experimental setups, like the number of total trials per condition and the asymmetry of number of trials between conditions. Additionally, we simulated different conditions by manipulating simulation parameters to affect PAC strength, noise strength and angle of opposition.

### 3.1. Testing distance measures sensitivity as a function of PAC and noise strength

In our first simulation, we explored the effect of PAC strength and the strength of pink noise. In our first simulation we calculated MOVI and JSD on a continuous scale of PAC strength and pink noise ranging from the no PAC to a strong PAC. **Figure 2** shows the number of significant hits over 100 simulated participants in case of opposed or non-opposed datasets.

**Figure 2.**
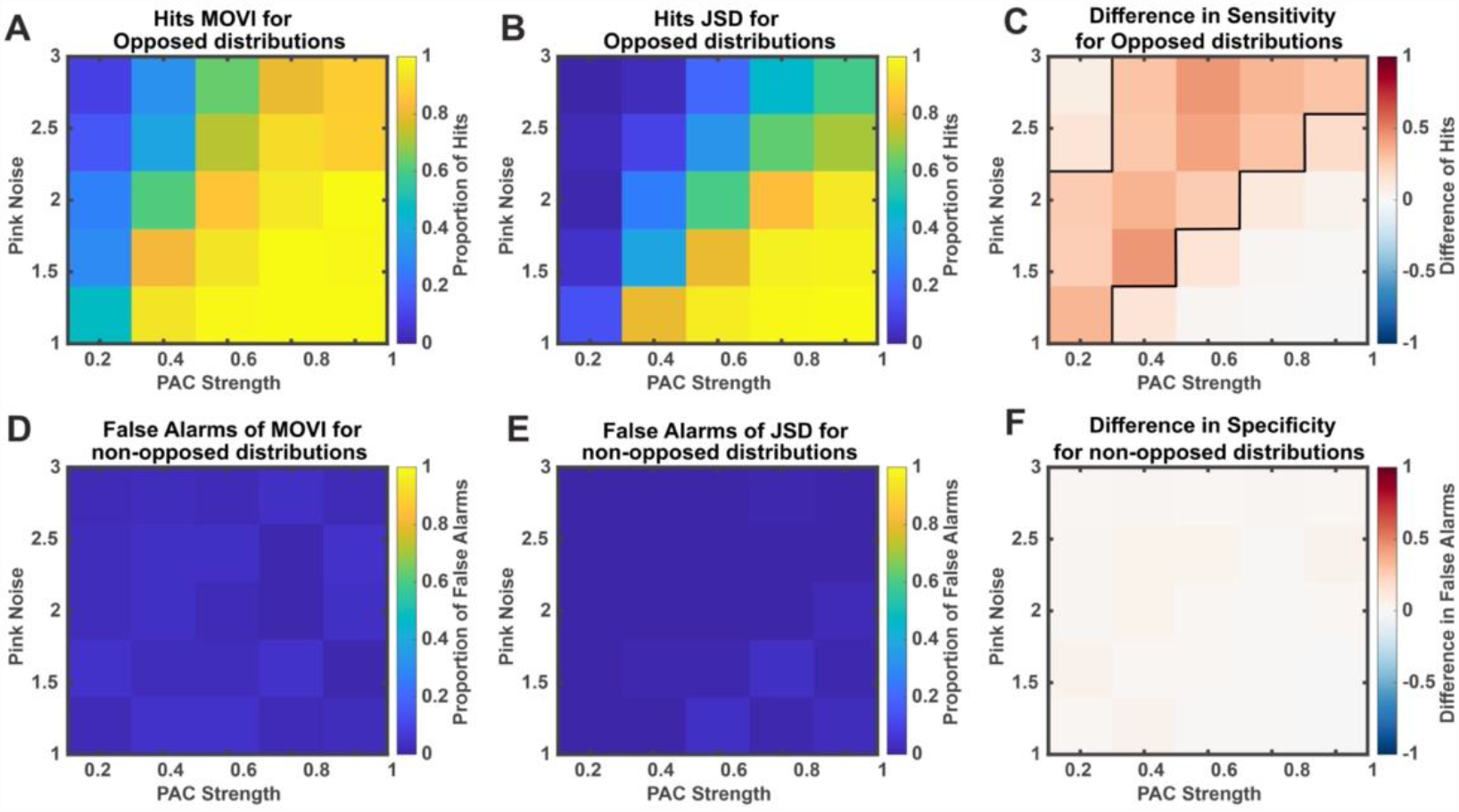
Hits and false alarms for MOVI and JSD in function of PAC and Pink Noise strength in Opposed and non-opposed distributions. (**A**) Proportion of Hits for MOVI on opposed distributions as a function of PAC strength and Pink Noise Strength. (**B**) Proportion of Hits for JSD on opposed distributions as a function of PAC strength and Pink Noise strength. (**C**) Difference in sensitivity between MOVI and JSD. A Binomial test was performed, and p-values were corrected with a Bonferroni correction. Threshold of p-values was a 0.05. For Opposed Distributions MOVI is significantly more sensitive than JSD for the settings within the cluster delimited by the dark line. (**D**) Proportion of False Alarms for MOVI for non-opposed distributions as a function of PAC strength and pink noise strength. (**E)**: Proportion of False Alarms for JSD for non-opposed distributions as a function of PAC strength and pink noise strength. (**F**): Difference in Specificity (1-False Alarms) between MOVI and JSD. No significant difference was detected between the two indexes.

With perfectly opposed distributions, MOVI is significantly more sensitive, and detects more easily opposed distributions than JSD. MOVI is capable of detecting opposition at lower levels of PAC compared to JSD.

We observed that both MOVI and JSD detected opposition with strong PAC in with low noise and opposition (**Figure 2A and B)** but not when the distributions have the same phase preference (**Figure 2D and E**). However, MOVI outperformed JSD to detect phase opposition at smaller PAC strength values and continued detecting phase opposition in conditions of larger noise values. MOVI exhibited a significantly higher sensitivity than JSD in those higher noise conditions as shown by the black line delimiting the cluster of values that survived correction depicted in **Figure 2C** (within cluster, binomial test, t_min_ = 3.49, p = 0.018, t_max_ = 6.04, p < 0.01). Furthermore, we calculated the average Positive Predictive Value of both indexes (see Methods) and across the matrices both indexes did not differ significantly in the average PPV (average PPV for MOVI = 0.94, average PPV for JSD = 0.97, t-test: t = -1.5, p = 0.11, df = 24). However, MOVI had a significantly higher NPV (MOVI NPV = 0.81, JSD NPV = 0.72, t-test: t = 7.24, p < 0.01, df = 24), thereby indicating it detected less false negatives than JSD. The accuracy of MOVI and JSD was 0.84 and 0.75 respectively, meaning that MOVI was more accurate in the detection of true positives and true negatives than JSD. The MCC of MOVI and JSD was 0.80 and 0.70 respectively. In these simulated conditions a result from MOVI is more likely to detect a real effect and is less biased than JSD.

### 3.2. Testing distance measures sensitivity as a function of angle difference and noise strength

In the previous test we showed that MOVI was more sensitive in identifying phase opposition compared to JSD with lower levels of PAC and at lower signal-to-noise ratio. Here, we examined whether these two methods differed in their sensitivity to phase opposition as a function of the degree of angle difference between two distributions and noise. To test this issue, we fixed PAC strength at a value of 0.4 where MOVI and JSD had a sensitivity and specificity that did not significantly differ (**Figure 2C and F**) and took angle difference as a variable. We generated various datasets by simulating angle differences in steps of 45 degrees, ranging from 0 to 180. Consequently, when the angle difference was 0, the distributions exhibited an identical phase preference, whereas with an angle difference of 180, the distributions displayed an opposite phase preference.

Results are depicted in **Figure 3**, where statistically significant differences (p < 0.05, corrected) between MOVI and JSD are outlined with a black line. Our results showed that MOVI was significantly better than JSD in detecting phase opposition, particularly when the angles of difference were smaller and in conditions of greater noise in the signals (within cluster, t_min_ = 3.39, p = 0.017, t_max_ = 5.25, p < 0.01, df = 24). Neither MOVI nor JSD detected phase opposition when there was none (**Figure 3A and B**), but JSD required less noisy data and greater angle difference between the distributions to detect phase opposition (**Figure 3C**). For higher noise values and phase preference differences starting at 90 degrees, MOVI exhibited greater sensitivity compared to JSD. In this test, we refrained from calculating the PPV, NPV, Accuracy, and MCC of the indexes. This decision was based on the difficulty of determining what constitutes a true positive or a true negative in the context of angle opposition variations. For instance, when the distance is 90 degrees, it is not possible to definitively classify the distributions as opposed or non-different.

**Figure 3.**
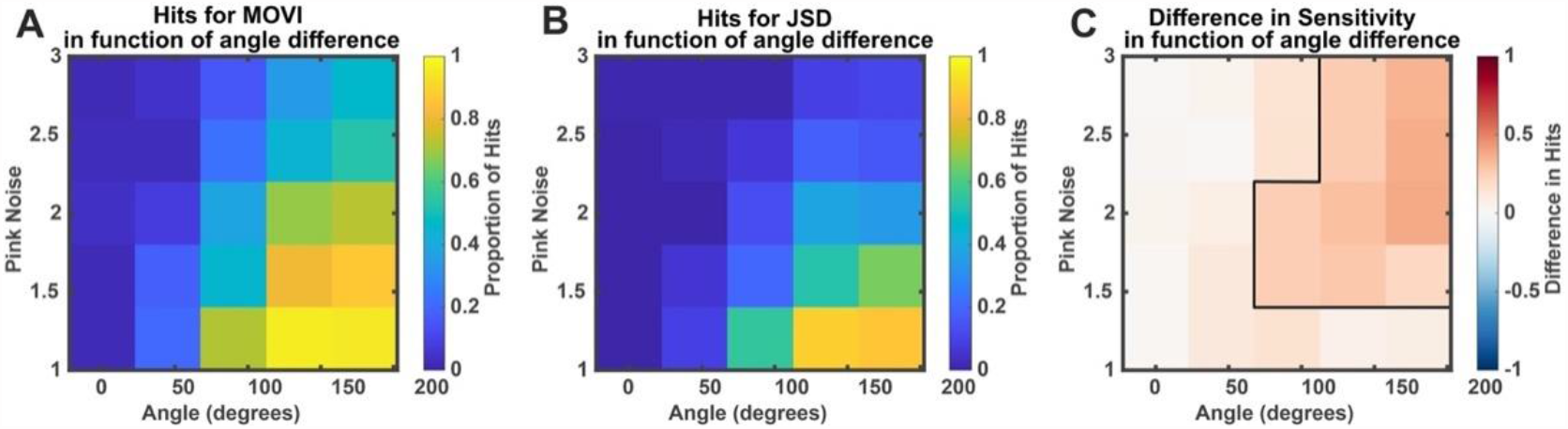
Hits for MOVI and JSD in function of angle difference between two distributions. (**A**) Proportion Of positive tests for MOVI as a function of angle difference between the two distributions and pink noise strength. (**B**) Proportion Of positive tests for JSD as a function of angle difference between the two distributions and pink noise strength. C: Difference in sensitivity between MOVI and JSD. A Binomial test was performed and p-values were corrected with a Bonferroni correction. Threshold of p-values was set at 0. For Opposed Distributions, MOVI is significantly more sensitive than JSD for the settings within the cluster delimited by the dark line that are higher noise strength and lower angle differences.

### 3.3. Are the number of trials important for MOVI and JSD?

We examined whether the sensitivity of MOVI and JSD changed as a function of number of trials. We simulated datasets with an increasing number of trials in both conditions (from 10 to 50 with increments of 10 trials) and calculated MOVI and JSD accordingly.

Results are depicted in **Figure 4**, where statistically significant differences at cluster level (p < 0.05, corrected) between MOVI and JSD are outlined with a black line. Our findings revealed that MOVI was capable of detecting phase opposition even with a smaller number of trials (**Figure 4A**), whereas JSD necessitated a higher number of trials in both conditions to detect phase opposition (**Figure 4B**). However, both indexes faced challenges in detecting phase opposition when both distributions consisted of only 10 trials. This implies that despite the superior sensitivity of MOVI compared to JSD, a substantial number of trials is still necessary to conduct the test effectively with MOVI. This observation could be attributed to the increased significance of factors like noise and random coupling between low and high frequencies, which are less likely to be mitigated through the computation of surrogate trials. Here too MOVI exhibits a significantly higher sensitivity than JSD (**Figure 4C**) as shown by the black line delimiting the cluster of values that survived correction (within cluster, t_min_ = 3.22, p = 0.031, t_max_ = 6.58, p < 0.01). In this case too, PPV did not differ significantly between MOVI and JSD (mean PPV for MOVI = 0.81, mean PPV for JSD = 0.87, t-test: t = -1.69, p = 0.10, df = 24). MOVI however had a significantly higher NPV than JSD (MOVI NPV = 0.57, JSD NPV = 0.52, t-test: t = 4.95, p < 0.01, df = 24), meaning that MOVI produced less false negatives (**Figure 4D, E and F**). Additionally, the Accuracy of MOVI was 0.61 and the accuracy of JSD was 0.53. The MCC of MOVI and JSD was respectively 0.40 and 0.21, showing that MOVI was more likely to detect a real effect than JSD.

**Figure 4.**
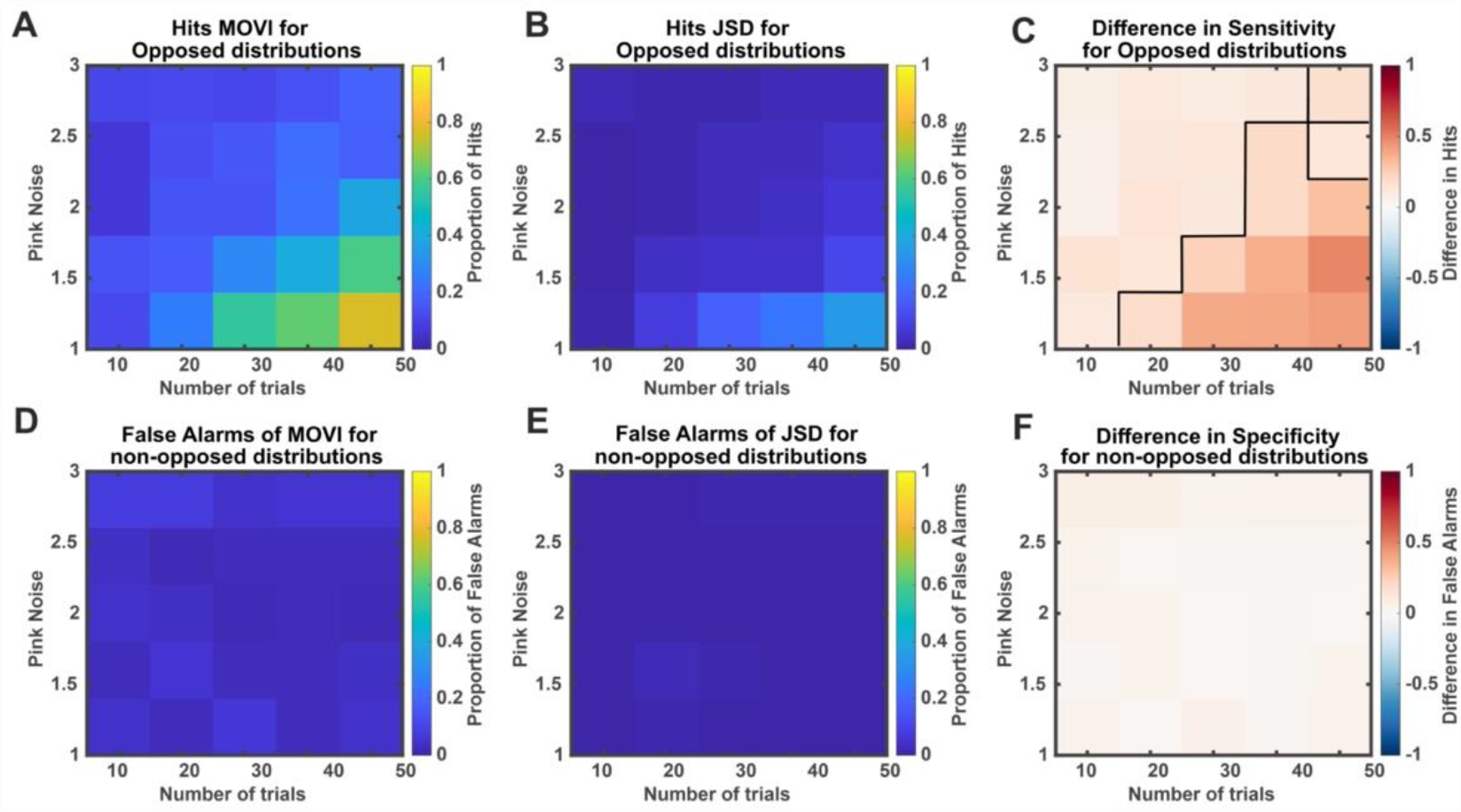
MOVI and JSD as a function of number of trials and noise. (**A**) Proportion of Hits for MOVI on opposed distributions as a function of number of trials and Pink Noise Strength. (**B**) Proportion of Hits for JSD on opposed distributions as a function of number of trials and Pink Noise strength. (**C**) Difference in sensitivity between MOVI and JSD. A binomial test was performed, and p-values were corrected with a Bonferroni correction. Threshold of p-values was a 0.05. For Opposed Distributions MOVI is significantly more sensitive than JSD for the settings within the cluster delimited by the dark line, where MOVI has a lower false omission rate even with a lower number of trials. (**D**) Proportion of False Alarms for MOVI for non-opposed distributions as a function of number of trials and pink noise strength. (**E**) Proportion of False Alarms for JSD for non-opposed distributions as a function of number of trials and pink noise strength. (**F**) Difference in Specificity (1-False Alarm Rate) between MOVI and JSD. No statistically significant difference was detected between the two indexes (all, p > 0.05).

### 3.4. Does asymmetrical number of trials between conditions influence MOVI and JSD?

In the previous test, we assessed both indexes using a symmetrical number of trials for each condition, ensuring that both conditions had an equal number of trials. However, we also wanted to investigate whether the indexes could detect opposition when there was an uneven number of trials between conditions. Furthermore, we sought to determine the extent to which this asymmetry could impact the sensitivity of both indexes. To this end, we varied the number of trials of only one distribution (between 10 and 50 with steps of 10) and the other distribution had a fixed number of 50 trials and a fixed PAC strength of 0.4.

Results are depicted in **Figure 5**, where statistically significant differences (p < 0.05, Bonferroni corrected) between MOVI and JSD are outlined with a black line (**Figure 5C**). What is observed is that it is generally not advised to work with a very low number of trials where false alarms appear in both MOVI and JSD for less than 20 trials in a single condition (**Figure 5D, E, and F)** as this could account for false positives. Overall, MOVI showed higher sensitivity than JSD with an uneven number of trials (within cluster, Binomial test, t_min_ = 3.48, p = 0.012, t_max_ = 6.52, p < 0.01). Through the calculation of the mean PPV, we found that a positive test result using JSD was more likely to be a true positive compared to MOVI (MOVI PPV = 0.85, JSD PPV = 0.92), and that this difference was statistically significant (t-test: t = - 3.31, p = 0.01, df = 24). Additionally, we computed the average NPV for both indexes (MOVI: NPV = 0.65; JSD: NPV = 0.56) and found that MOVI was significantly less prone to false negatives (t-test: t = 8.27, p < 0.01, df = 24). we obtained an Accuracy value of 0.70 for MOVI and 0.59 for JSD, while their MCC were 0.57 and 0.41, respectively. Once again, in this test, MOVI demonstrated a greater likelihood of detecting genuine effects compared to JSD.

**Figure 5.**
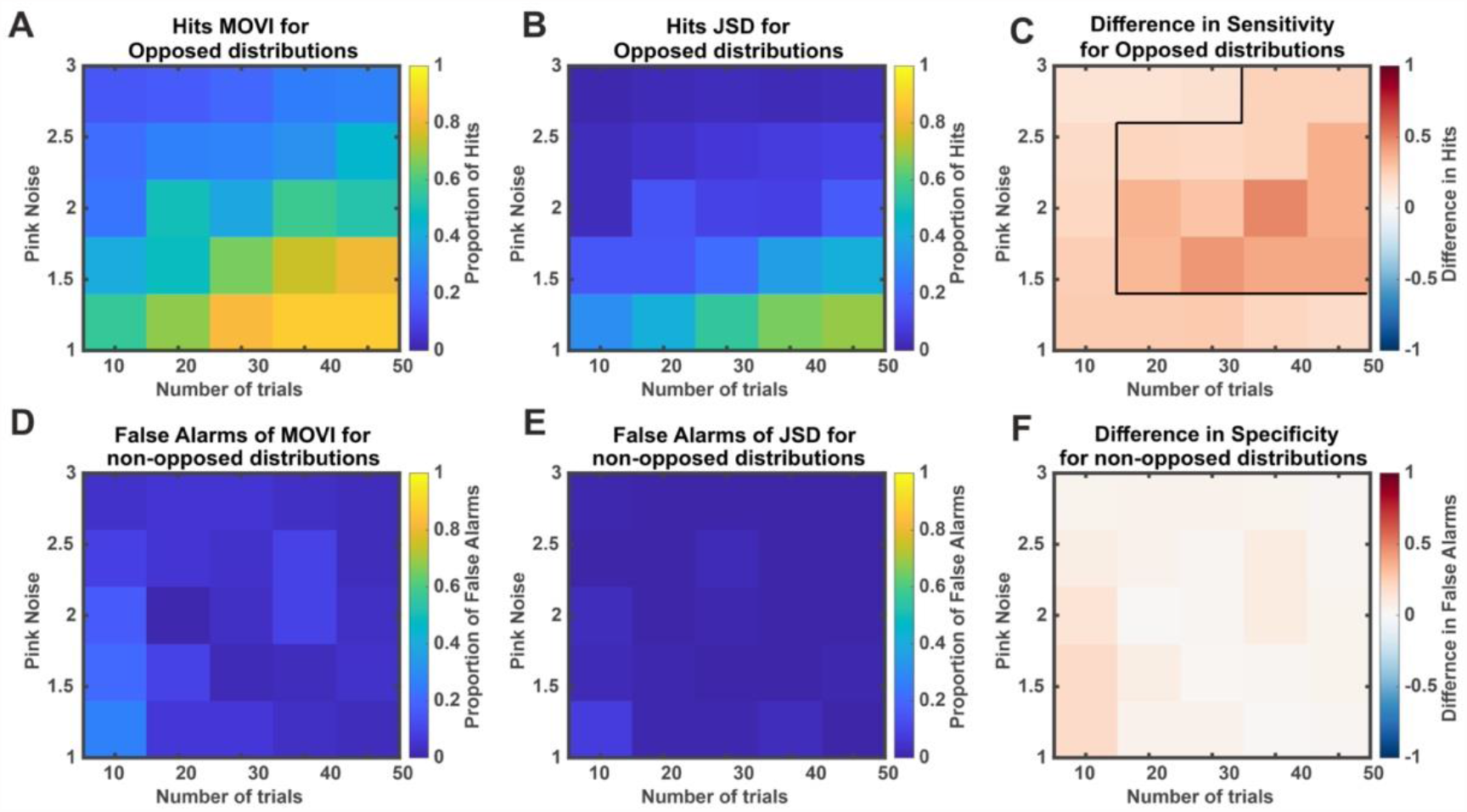
MOVI and JSD as a function of asymmetrical number of trials. (**A**) Proportion of Hits for MOVI on opposed distributions as a function of number of trials in only one distribution (asymmetrical number of trials) and Pink Noise Strength. (**B**) Proportion of Hits for JSD on opposed distributions as a function of number of trials for one distribution and Pink Noise strength. (**C**) Difference in sensitivity between MOVI and JSD. A Binomial test was performed, and p-values were corrected with a Bonferroni correction. Threshold of p-values was set at 0.05. For Opposed Distributions, MOVI was significantly more sensitive than JSD for the settings within the cluster delimited by the dark line, where MOVI has a lower false omission rate even with a lower number of trials in only one condition. (**D**) Proportion of False Alarms for MOVI for non-opposed distributions as a function of number of trials and pink noise strength. (**E**) Proportion of False Alarms for JSD for non-opposed distributions as a function of number of trials and pink noise strength. (**F**) Difference in Specificity (1-False Alarm Rate) between MOVI and JSD. No significant difference was detected between the two indexes, although one value almost survived Bonferroni correction with pink noise = 1 and 10 trials in only one condition (binomial test: t = 3.32, p = 0.063).

### 3.5. Does inter-trial coherence affect MOVI and JSD?

Here, while keeping the Phase-Amplitude Coupling (PAC) strength fixed, we assessed whether different degrees of coherence in phase preference for each individual trial influenced MOVI and JSD indexes. Thus, in this analysis, every trial is expected to display PAC, albeit with varying levels of inter-trial phase preference. Consequently, in scenarios where there is a high variability in inter-trial phase preference, measuring the coupling strength of each individual trial would likely yield significant coupling. However, if we were to average the distributions of all trials and then measure the coupling strength of the average, it is probable that we would not observe significant coupling. This distinction is relevant, for example, in cases where experimenters preferred to test PAC on each trial individually (Axmacher et al., 2010).

Results are depicted in **Figure 6**, where statistically significant differences at cluster level (p < 0.05, corrected) between MOVI and JSD are outlined with a black line. Our findings revealed that both MOVI and JSD detected shared and consistent differences in phase preference (**Figure 6A, B, D, and E)** However, MOVI exhibited greater sensitivity in cases of higher inter-trial variability in angle preference and higher levels of pink noise (**Figure 6C**) (within cluster, binomial test, t_min_ = 3.20, p = 0.034, t_max_ = 5.84, p < 0.01). We also found that although MOVI showed slightly more false positives this decrease in specificity was not significant compared to JSD (**Figure 6F**). Additionally, we calculated the average PPV of both indexes (PPV for MOVI = 0.90, PPV for JSD = 0.96) and compared them. This analysis determined that a positive test result with JSD was more likely to represent a true positive compared to MOVI under these simulation conditions (t-test, t= -3.75, p = 0.01, df = 24).

**Figure 6.**
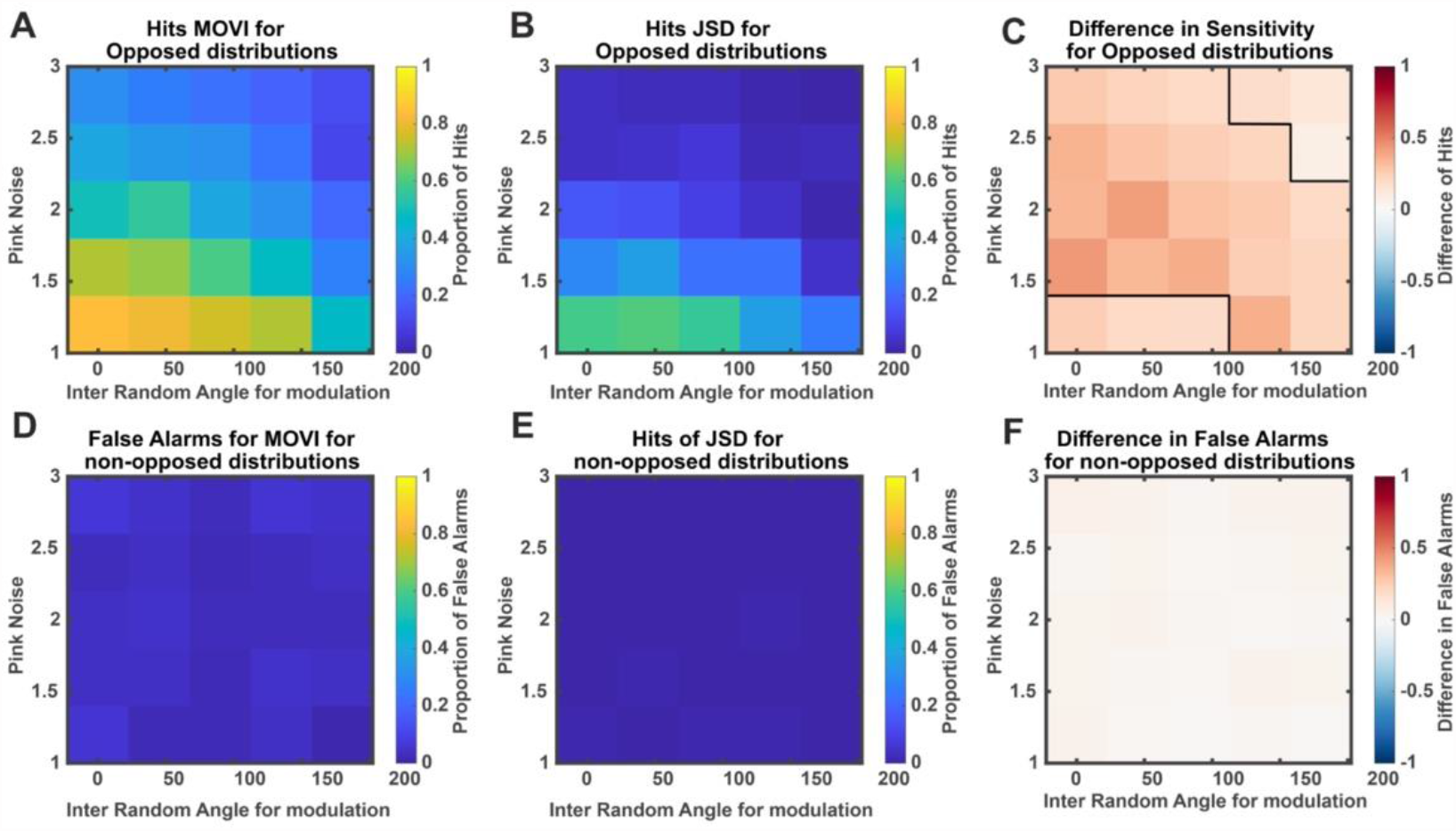
MOVI and JSD as a function of Inter-Trial phase-preference variability. (**A**) Proportion of Hits for MOVI on opposed distributions as a function of Inter-trial coherence of angle preference and Pink Noise Strength. (**B**) Proportion of Hits for JSD on opposed distributions as a function of Inter-trial coherence of angle preference and Pink Noise strength. (**C**) Difference in sensitivity between MOVI and JSD. A binomial test was performed, and p-values were corrected with a Bonferroni correction. Threshold of p-values was a 0.05. For Opposed Distributions MOVI is significantly more sensitive than JSD for the settings within the cluster delimited by the dark line, where MOVI has a lower false omission rate with higher inter-trial variability in angle preference. (**D**) Proportion of False Alarms for MOVI for non-opposed distributions as a function of Inter-trial coherence of angle preference and pink noise strength. (**E**) Proportion of False Alarms for JSD for non-opposed distributions as a function of Inter-trial coherence of angle preference and pink noise strength. (**F**) Difference in Specificity (1-False Alarm Rate) between MOVI and JSD. No significant difference was detected between the two indexes.

We computed the NPV for both indexes, resulting in an NPV of 0.65 for MOVI and 0.55 for JSD. A direct comparison of the two showed that a negative result by MOVI was significantly (t-test, t = 9.65, p < 0.01, df =24) more likely to be truly negative, while JSD exhibited a significantly higher rate of false negatives. Average Accuracy for MOVI was 0.70 and for JSD 0.59. The MCC for MOVI was 0.59, and for JSD it was 0.40. In sum, these results demonstrate that MOVI displayed superior accuracy in detecting real positive or negative effects and produced fewer false results compared to JSD.

## 4. Discussion

Phase-Amplitude Coupling (PAC) is widely studied phenomenon in the brain, and its mechanisms offer insights into how the brain processes and represents information. Computational models propose that nested gamma oscillations within theta cycles may organize sequences of events, while other models suggest gamma couples to opposed-phase states during memory encoding and retrieval. Characterizing differences in phase preference in PAC analysis between neural states or experimental conditions is crucial for understanding neural coding and mnemonic operations. In the current study, we presented MOVI, an analysis method for detecting phase preference difference within PAC between two datasets. Using synthetic data, we showed that at MOVI is more tolerant to low signal-to-noise ratio, lower levels of angle differences and trial number asymmetry between conditions than other robust phase-opposition indexes like JSD.

We found that MOVI is more sensitive than JSD in detecting phase opposition. MOVI can detect phase differences even when PAC is weaker, with fewer or unevenly distributed trials, and smaller angle variations between distributions. It is also more resilient to noise and variations in angle preference across trials. While this higher sensitivity may lead to more false negatives, it does not significantly impact the overall accuracy of the Matthews Correlation Coefficient (MCC), a measure of performance in binary classification tasks. In fact, in all tests conducted, MOVI consistently outperformed JSD in terms of accuracy and MCC, demonstrating its superior sensitivity to real differences and its ability to effectively detect true positive and negative effects, primarily by reducing the number of false negatives compared to JSD. These findings highlight the advantages of using MOVI over JSD for quantifying phase opposition in neurophysiological data, particularly when dealing with complex and noisy signals. MOVI’s ability to detect smaller phase differences and handle varying trial conditions makes it a valuable tool for understanding phase-preference patterns and phase-amplitude coupling in the brain.

While MOVI proves highly effective in detecting phase opposition and capturing phase-amplitude coupling patterns, it may not be the ideal choice for analyzing event occurrences like spiking frequency (Rutishauser, 2019; Rutishauser et al., 2010) or ripple counts (Norman et al., 2019) These scenarios involve instances that do not necessarily follow a probabilistic distribution shape, and circular statistics (Berens, 2009) PPC (Vinck, van Wingerden, Womelsdorf, Fries, & Pennartz, 2010), ITC (VanRullen, 2016), or PACOI (Costa et al., 2022) might offer more suitable alternatives. These methods consider different aspects of neural activity and are better suited to address specific research questions where the distribution shape might not be appropriate for MOVI analysis.

However, one key advantage of MOVI compared to other methods is its adaptability to studying phase-amplitude phenomena without the need to summarize the power dimension of amplitude as unit-normalized vectors. MOVI and JSD both allow the treatment of amplitude data without any loss of information by modifying the neurophysiological data into another form. However, MOVI stands out as being more sensitive and capable of accurately discriminating phase-preference opposition or its absence within PAC between conditions.

Another beneficial aspect of MOVI is that it generates an index of opposition for each observation or subject. This feature facilitates a smoother transition to second-level statistics, where experimental MOVI values can be compared to surrogate MOVI values. Consequently, MOVI can serve as an effective exploratory method in the study of phase opposition within PAC distributions, enabling researchers to investigate previously unknown interactions between mechanisms in neurophysiology and experimental settings or neural states.

One limitation of our study is that we evaluated the MOVI and JSD methods using a 2.5s segment of artificial data. While this segment is suitable for most task-induced research in humans and animals, it may not fully capture the dynamics of neural activity during extended periods or other types of research focusing on phase-preference within PAC over longer time frames. For instance, in animal studies, researchers often record neural activity over extended periods to capture a comprehensive view of brain dynamics while navigating in a maze or during states of quiescence. Similarly, during sleep studies, neural data is typically recorded over hours to observe the brain’s activity during different sleep stages. Nevertheless, while our study did not specifically test MOVI for long time segments, we believe that MOVI, given its sensitivity to phase-preference opposition and its adaptability to various data lengths, would still be a valid approach to assess phase preference differences in such scenarios.

In conclusion, MOVI stands out as a powerful and versatile tool in the neuroscientific field. Its unique ability to combine coupling strength and opposition measures makes it an attractive choice for analyzing complex neural interactions and mechanisms in neurophysiology signal datasets. Unlike other indexes, MOVI does not necessitate additional verification of coupling strength, allowing for a more straightforward exploration of phase-preference differences between datasets. Furthermore, the user-friendly nature of MOVI makes it intuitive and easy to use, providing a comprehensive summary of unidirectional phase-preference differences between datasets. Overall, MOVI’s capabilities make it a valuable addition to the neuroscientific toolbox, facilitating the exploration of phase-preference patterns and the investigation of neural dynamics in diverse experimental settings.

## 5. Conclusions

In this paper, we introduced a novel method to analyze phase-preference variations within PAC in neurophysiological data: the Mean Opposition Vector Length Index. Our approach involved creating an alternate distribution and measuring its non-uniformity using the Mean Vector Length method. Through synthetic experiments, we demonstrated that MOVI showed superior sensitivity compared to a conventional measure (JSD) in various settings. We suggest that MOVI offers greater flexibility and adaptability to neurophysiological data and practical experiments. It proved to be more robust against low number of trials, noise, and asymmetrical numbers of trials that commonly occur in experimental conditions and neurophysiological recordings. We conclude that MOVI is well-suited for real-world research scenarios, enabling more accurate and comprehensive analysis of phase-preference differences within PAC patterns between diverse neurophysiological datasets.

## Funding

This work was supported by the Spanish Ministerio de Ciencia, Innovación y Universidades, which is part of Agencia Estatal de Investigación (AEI), through the project PID2019-111199GB-I00 (Co-funded by European Regional Development Fund. ERDF, a way to build Europe), to L.F and the Programa de retos: Jóvenes Investigadores: JIN RTI2018-100977-J-I00 awarded to APB. We thank CERCA Programme/Generalitat de Catalunya for institutional support.

### CRediT authorship contribution statement

**Ludovico Saint-Amour di Chanaz:** Conceptualization, Methodology, Data curation, Writing – original draft, Visualization, Investigation, Validation, Writing – review & editing.

**Alexis Pérez-Bellido:** Conceptualization, Methodology, Visualization, Writing – review & editing.

**Lluís Fuentemilla:** Conceptualization, Visualization, Writing – review & editing.

### Data Availability

The data and codes that support the findings of this study are available on GitHub https://github.com/DMFresearchlab/MOVICode

## Acknowledgments

We thank Daniel Pacheco-Estefan and Diego Lozano-Soldevilla for their helpful discussions on earlier versions of the manuscript.

